# Unsupervised tensor integration of hippocampal and colonic multi-omics reveals enteric neural signatures in cognitive aging

**DOI:** 10.64898/2026.11.20.768248

**Authors:** Y-H Taguchi, Turki Turki

## Abstract

**Background:** Molecular changes in peripheral tissues may provide insights into cognitive aging, but identifying relevant genes without using behavioral out-comes for feature selection remains challenging.

**Results:** We applied tensor decomposition-based unsupervised feature extraction to publicly available hip-pocampal and colonic transcriptomic and promoter DNA methylation profiles from mice across five age groups. A tensor comprising tissue, molecular modality, age group, biological replicate, and gene dimensions was constructed using 16,080 shared genes. Higher-order singular value decomposition identified a component with a monotonic age pattern, shared tissue contributions, and opposing transcriptomic and methylation contributions. This component yielded 1,577 selected genes without using cognitive scores for feature selection. The corresponding colonic methylation score correlated with overall cognition (***r* = 0.589**; Benjamini–Hochberg-adjusted ***P* = 9.87 *×* 10**^***−*7**^), although no significant associations remained after age adjustment. The selected genes were enriched for enteric neuronal and glial markers derived from a published single-cell atlas, with 5.10-fold and 4.65-fold enrichment, respectively. Comparison with the broadest available Boruta-selected gene set identified only 64 overlapping genes, leaving 1,513 genes selected exclusively by the tensor approach in this comparison. These remaining genes retained significant enrichment for synaptic processes and enteric neuronal and glial signatures using library-specific backgrounds.

**Conclusions:** Unsupervised tensor integration recovered an aging-associated relationship between colonic methylation and cognition while identifying a largely distinct set of functionally coherent candidate genes. These findings complement supervised feature selection and provide candidates for investigating enteric neural involvement in cognitive aging, without establishing an age-independent association or a causal gut-to-brain mechanism.

## 1 Background

Our ability to perceive the world, remember past experiences, and form thoughts depends critically on the brain. The profound changes in cognition and awareness that can follow brain injury make it natural to regard the brain as the central organ of mental life. Yet the importance of the brain does not mean that its functions can be understood in isolation from the body. The brain continuously receives signals from peripheral organs, and these signals can influence how we perceive, feel, and behave. Understanding cognition therefore requires consideration not only of processes within the brain, but also of its interactions with the rest of the body.

The skin provides an illustrative example. It is not merely a protective barrier or a passive surface for receiving external stimuli. Human hair follicles possess a functional equivalent of the hypothalamic–pituitary–adrenal axis and can synthesize cortisol, demonstrating neuroendocrine activity outside the brain [1]. Furthermore, gentle touch conveyed by unmyelinated tactile afferents is associated with insular cortex activation and affective aspects of touch [2]. Thus, the skin illustrates how peripheral sensory and endocrine functions can contribute to processes relevant to emotional experience.

The heart provides another example. Although its association with emotion has traditionally been expressed metaphorically, experimental evidence shows that cardio-vascular signals can influence emotional perception. In humans, fearful faces presented during cardiac systole were detected more readily and judged as more intense than those presented during diastole. These effects were accompanied by stronger amyg-dala responses [3]. The relationship between the heart and emotion therefore has a physiological basis: signals associated with the heartbeat can modulate how the brain processes emotionally salient information.

Among these interactions between peripheral organs and the brain, the gut–brain axis has attracted particular interest. The gut provides a setting in which host tissues, the enteric nervous system, and resident microorganisms may all contribute to influences on brain function. Experimental manipulation of the intestinal microbiota, including transfer between mouse strains, has been shown to alter exploratory behavior and brain-derived neurotrophic factor levels [4]. More specifically, supplementation with *Lactobacillus reuteri* restored oxytocin-related measures, social interaction-induced synaptic plasticity, and social behavior in offspring of mice fed a maternal high-fat diet [5]. Such findings have motivated the concept of psychobiotics, which refers to live microorganisms with potential mental health benefits [6]. They demonstrate that influences arising from the gut can have measurable consequences for brain function and behavior.

These observations are also relevant to cognitive aging. Transplantation of microbiota from young donors into aged mice attenuated selected age-associated cognitive deficits and altered hippocampal transcriptomic and metabolomic profiles [7]. However, these experimental findings do not fully explain which host molecular changes in the gut and brain accompany cognitive aging. An important next step is therefore to investigate molecular profiles in both tissues within a common analytical framework. Integrating gene expression and DNA methylation profiles offers an opportunity to identify shared age-related patterns and candidate genes associated with the gut–brain axis.

We previously [8] analyzed gene expression and miRNA profiles of various organs including brains with tensor decomposition based unsupervised feature extraction [9] and found that PTSD induced coexpression of genes and miRNAs between brain and heart. Flor et al. [10] reported an association between colonic DNA methylation and cognitive performance during aging. The purpose of the present study is to reanalyze these datasets using TD-based unsupervised feature extraction to identify genes and biological processes associated with cognitive aging without using cognitive scores for gene selection.

## 2 Results

### 2.1 Identification of a shared age-related molecular component

At first, we need to identify 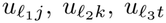, and 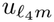. As expected, 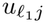 and 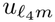 are independent of *j* and *m*, respectively, 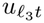 is monotonic, and 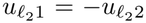 when (*ℓ*_1_, *ℓ*_2_, *ℓ*_3_, *ℓ*_4_) = (1, 1, 2, 1)(Fig. 1).

**Fig 1.**
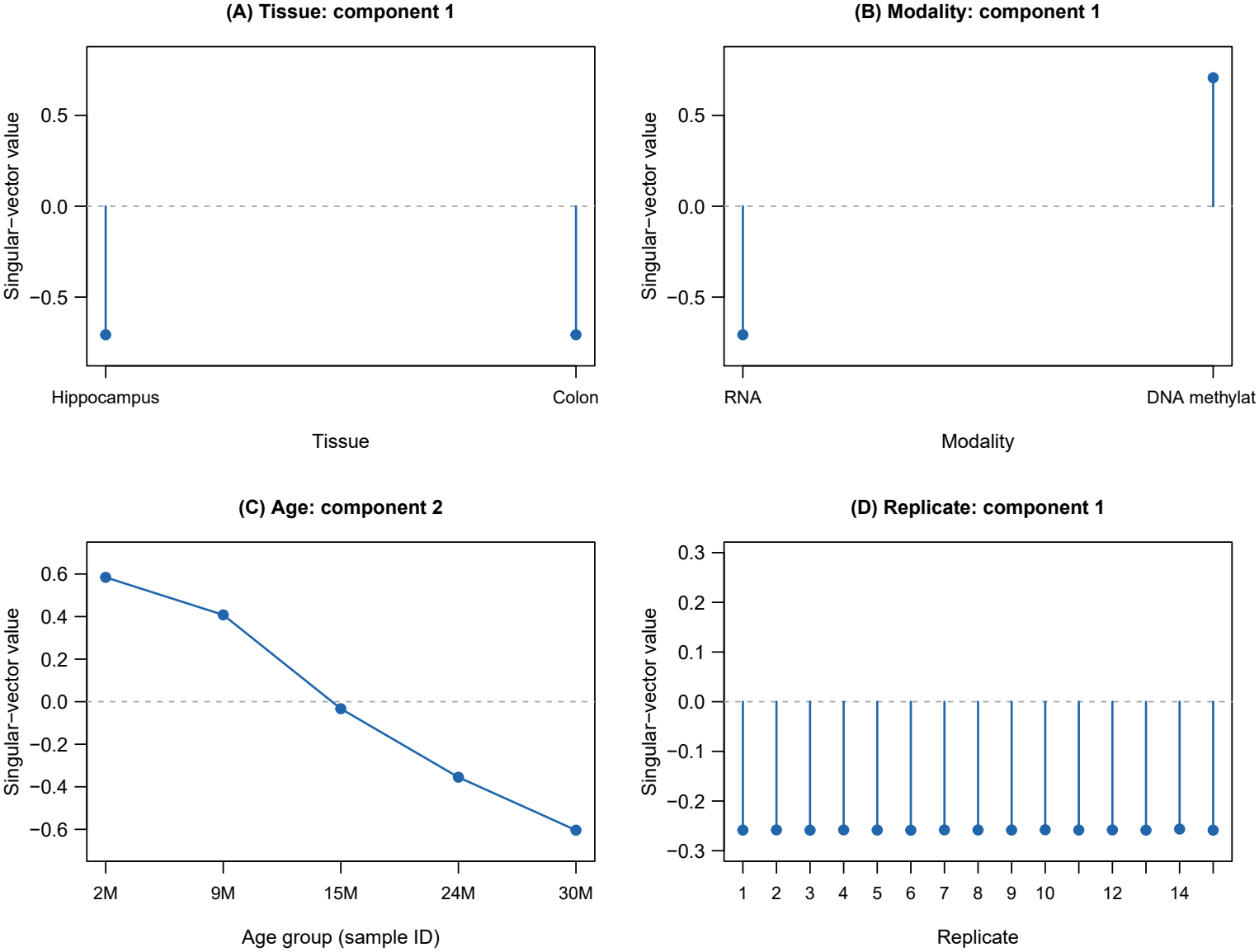
Selected mode-specific singular vectors from the HOSVD of the gene-level tensor. (A) The first tissue-mode singular vector. (B) The first modality-mode singular vector. (C) The second age-mode singular vector, with age groups labeled using the original sample-ID prefixes. The labels 2M, 9M, 15M, 24M, and 30M correspond to the study’s reported ages of 3, 9, 15, 24, and 28 months, respectively. (D) The first replicate-mode singular vector. The tensor comprises tissue × modality × age group × replicate × gene. Each vector has unit Euclidean norm and is shown without singular-value weighting. Dashed horizontal lines indicate zero. Replicate indices represent positions within each age group and do not identify the same animal across ages.

### 2.2 Gene selection from the associated gene-mode component

Next we need to specify *ℓ*_5_ associated with the absolute largest *G*(1, 1, 2, 1, *ℓ*_5_) (Fig. 2). It is obvious that *ℓ*_5_ = 5 is associated with the absolute largest *G*(1, 1, 2, 1, *ℓ*_5_). Thus, we decide to use *u*_5*i*_ to select genes, *i*s (Fig. 3). As a result, we selected 1,577 genes.

**Fig 2.**
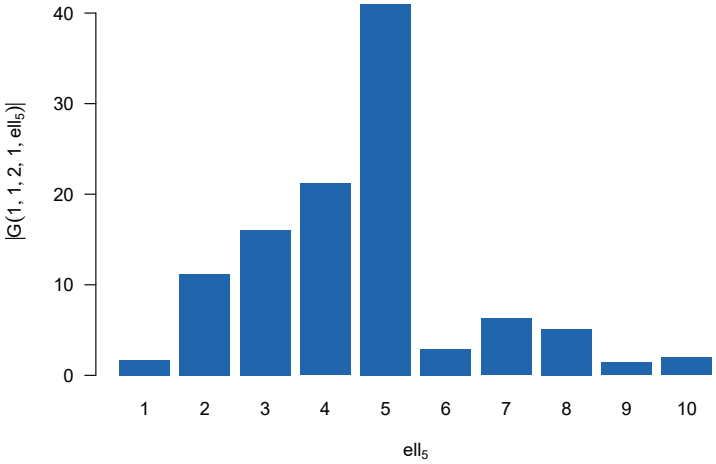
Absolute core-tensor coefficients for the first ten gene-mode components. Bars show |*G*(1, 1, 2, 1, *ℓ*_5_)| for *ℓ*_5_ = 1, …, 10, with the tissue, modality, age, and replicate component indices fixed at 1, 1, 2, and 1, respectively.

**Fig 3.**
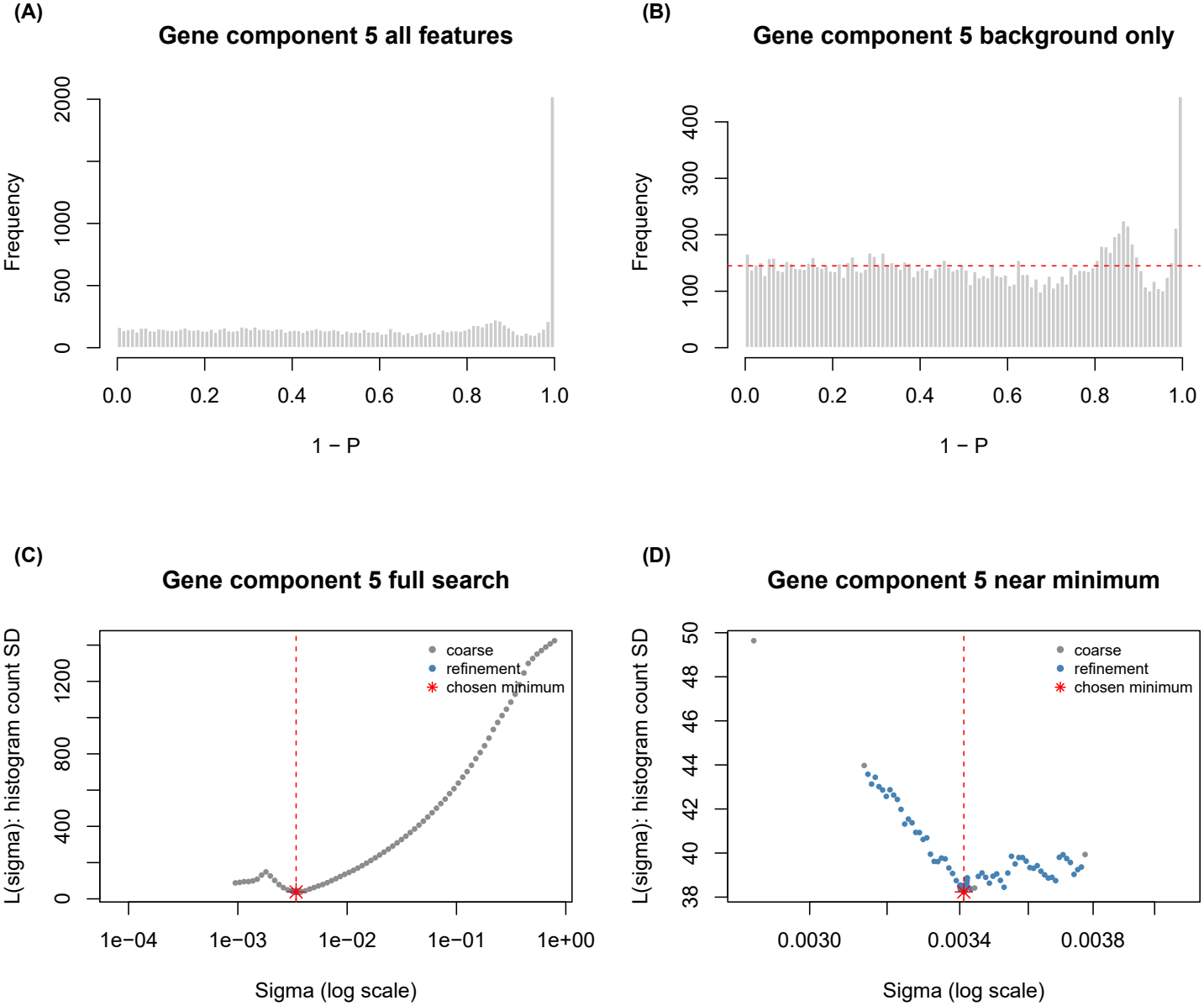
Diagnostics of standard deviation optimization for gene selection using the fifth gene-mode singular vector. (A) Distribution of 1 *− P* for all genes at the optimized standard deviation. (B) Corresponding distribution for the background genes retained during optimization; the red horizontal line indicates the mean bin count. (C) Objective function *L*(*σ*), defined as the standard deviation of histogram bin counts for the background genes, over the full search range. (D) Enlarged view near the selected minimum. Gray and blue points represent the coarse and refinement searches, respectively. Red asterisks and vertical dashed lines indicate the selected minimum and corresponding *σ*. The horizontal axes in (C) and (D) are logarithmic. P-values were calculated from the mean-centered gene loadings under a Gaussian assumption, and genes with Benjamini–Hochberg-adjusted P-values *≤* 0.01 were selected.

### 2.3 Enrichment analysis

We uploaded a set of 1,577 genes to Enrichr; before submission to Enrichr, Ensembl mouse gene identifiers (ENSMUSG IDs) were converted to gene symbols using the Ensembl release 116 annotation for the GRCm39 mouse genome assembly (Mus_musculus.GRCm39.116.gtf.gz). Gene identifier–symbol mappings were extracted from the gene id and gene name attributes of gene records in the GTF file. Version suffixes were removed from the input identifiers before matching. Unmapped identifiers and identifiers associated with multiple distinct symbols were flagged and excluded from the enrichment input. The resulting gene symbols were deduplicated, yielding 1,574 unique symbols for submission to the standard Enrichr service. The full list of 1,577 selected Ensembl gene identifiers and their symbol mappings is provided in Additional file 1.

The complete archived Enrichr outputs are provided in Additional file 2. For presentation, we highlight terms relevant to aging and neural functions; these examples do not constitute a separate confirmatory test of a pre-specified gut–brain hypothesis. Table 1 shows the top ten enriched terms in “Orphanet Augmented 2021” library in Enrichr. It is obvious that the set of genes includes many neuronal disease related genes, thus it is reasonable. Table 2 lists the top ten terms in “GTEx Aging Signatures 2021” library in Enrichr. Since it detects the association with aging, it is also reasonable. It should also be noted that SmallIntestine is listed although colon itself is not listed (colon is ranked in the twelfth with the adjusted *P*-value 0.028).

**Table 1.**
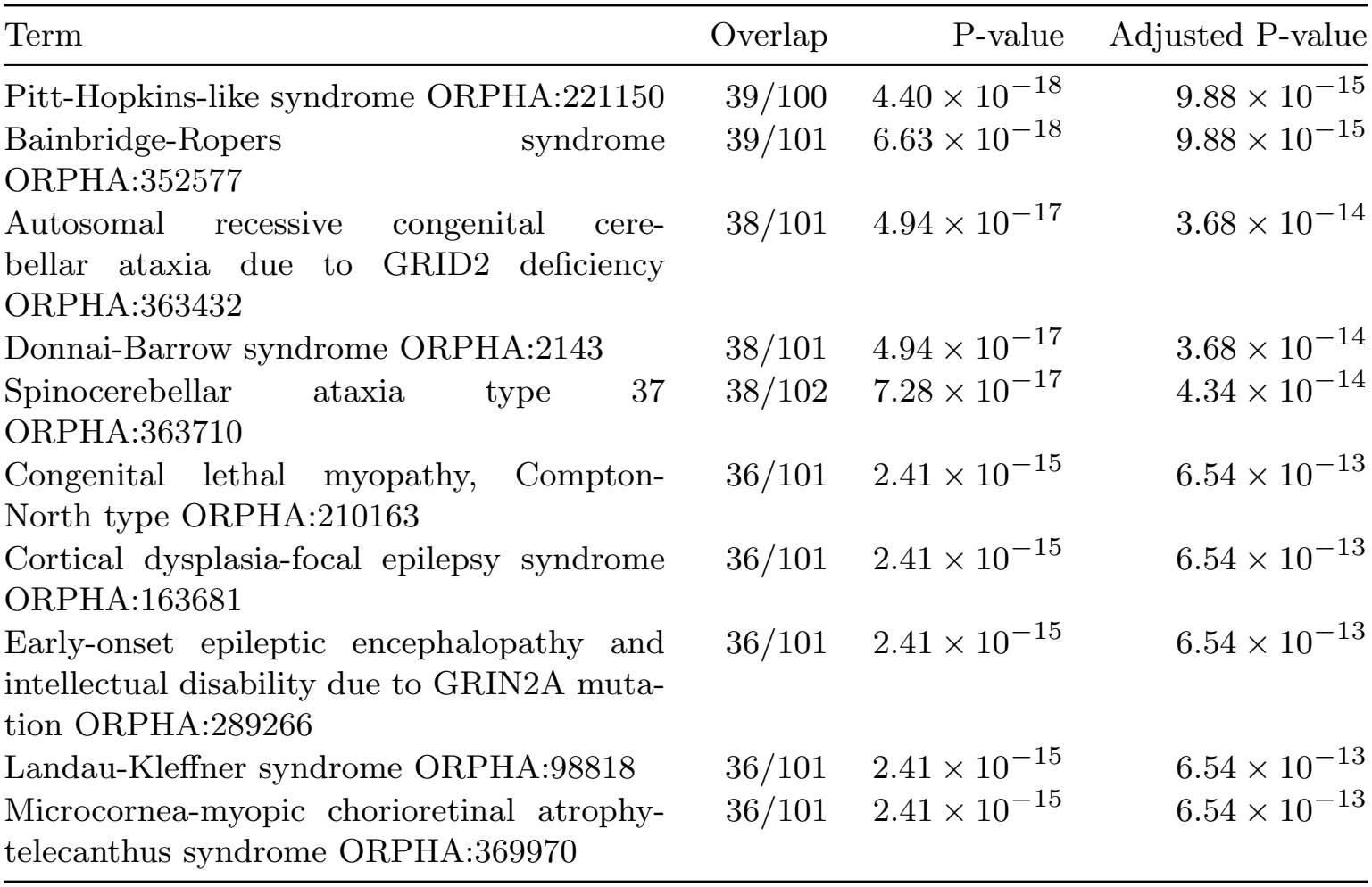
Top 10 enrichment results from the Orphanet Augmented 2021 library in Enrichr, shown in the order of the saved output. Overlap denotes the number of input genes shared with each gene set divided by the gene-set size.

| Term | Overlap | P-value | Adjusted P-value |
|---|---|---|---|
| Pitt-Hopkins-like syndrome ORPHA:221150 | 39/100 | 4.40 × 10<sup>−18</sup> | 9.88 × 10<sup>−15</sup> |
| Bainbridge-Ropers syndrome ORPHA:352577 | 39/101 | 6.63 × 10<sup>−18</sup> | 9.88 × 10<sup>−15</sup> |
| Autosomal recessive congenital cere-bellar ataxia due to GRID2 deficiency ORPHA:363432 | 38/101 | 4.94 × 10<sup>−17</sup> | 3.68 × 10<sup>−14</sup> |
| Donnai-Barrow syndrome ORPHA:2143 | 38/101 | 4.94 × 10<sup>−17</sup> | 3.68 × 10<sup>−14</sup> |
| Spinocerebellar ataxia type 37 ORPHA:363710 | 38/102 | 7.28 × 10<sup>−17</sup> | 4.34 × 10<sup>−14</sup> |
| Congenital lethal myopathy, Compton-North type ORPHA:210163 | 36/101 | 2.41 × 10<sup>−15</sup> | 6.54 × 10<sup>−13</sup> |
| Cortical dysplasia-focal epilepsy syndrome ORPHA:163681 | 36/101 | 2.41 × 10<sup>−15</sup> | 6.54 × 10<sup>−13</sup> |
| Early-onset epileptic encephalopathy and intellectual disability due to GRIN2A muta-tion ORPHA:289266 | 36/101 | 2.41 × 10<sup>−15</sup> | 6.54 × 10<sup>−13</sup> |
| Landau-Kleffner syndrome ORPHA:98818 | 36/101 | 2.41 × 10<sup>−15</sup> | 6.54 × 10<sup>−13</sup> |
| Microcornea-myopic chorioretinal atrophy-telecanthus syndrome ORPHA:369970 | 36/101 | 2.41 × 10<sup>−15</sup> | 6.54 × 10<sup>−13</sup> |

**Table 2.** Top 10 enrichment results from the GTEx Aging Signatures 2021 library in Enrichr, shown in the order of the saved output. Overlap denotes the number of input genes shared with each gene set divided by the gene-set size. Term names, including Up and Down labels, are retained as provided by Enrichr.

| Term | Overlap | P-value | Adjusted P-value |
|---|---|---|---|
| GTEx SmallIntestine 20-29 vs 50-59 Up | 47/250 | 1.77 × 10<sup>−8</sup> | 4.77 × 10<sup>−6</sup> |
| GTEx Brain 20-29 vs 60-69 Down | 45/250 | 1.34 × 10<sup>−7</sup> | 1.81 × 10<sup>−5</sup> |
| GTEx Nerve 20-29 vs 70-79 Up | 42/250 | 2.28 × 10<sup>−6</sup> | 2.05 × 10<sup>−4</sup> |
| GTEx Nerve 20-29 vs 50-59 Up | 37/250 | 1.46 × 10<sup>−4</sup> | 6.56 × 10<sup>−3</sup> |
| GTEx SmallIntestine 20-29 vs 40-49 Up | 37/250 | 1.46 × 10<sup>−4</sup> | 6.56 × 10<sup>−3</sup> |
| GTEx Ovary 20-29 vs 50-59 Up | 37/250 | 1.46 × 10<sup>−4</sup> | 6.56 × 10<sup>−3</sup> |
| GTEx Brain 20-29 vs 70-79 Down | 36/250 | 3.07 × 10<sup>−4</sup> | 1.18 × 10<sup>−2</sup> |
| GTEx Nerve 20-29 vs 60-69 Up | 35/250 | 6.25 × 10<sup>−4</sup> | 2.11 × 10<sup>−2</sup> |
| GTEx Ovary 20-29 vs 60-69 Up | 34/250 | 1.24 × 10<sup>−3</sup> | 2.78 × 10<sup>−2</sup> |
| GTEx Muscle 20-29 vs 60-69 Up | 34/250 | 1.24 × 10<sup>−3</sup> | 2.78 × 10<sup>−2</sup> |

### 2.4 Correlation with cognition score

Although these are only two examples that demonstrate that the selected set of genes are related to both cognition and aging, it will be better to relate *u*_5*i*_ to individual cognition score. We have computed the correlation between *x*_*jktm*5_ and *c*_*jktm*_ (Table 3). Most of the significant correlations were observed between colonic methylation and cognitive scores (Table 3).

**Table 3.**
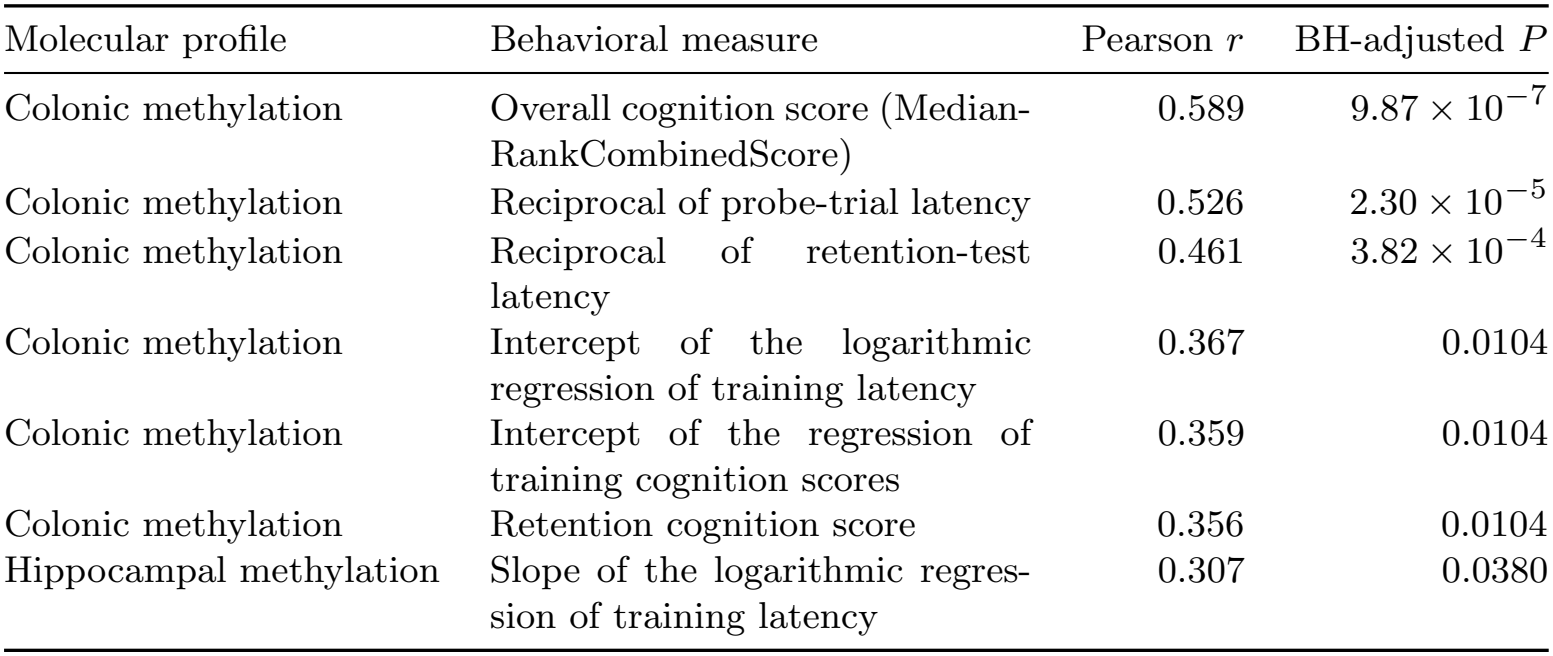
Significant correlations between fifth-component molecular scores and behavioral measures. Molecular scores were obtained by projecting normalized gene profiles onto the fifth gene-mode singular vector. Pearson correlations were calculated across 75 mice without age adjustment. P-values were adjusted using the Benjamini–Hochberg (BH) procedure across all 36 tests. Only associations with BH-adjusted P-values below 0.05 are shown.

| Molecular profile | Behavioral measure | Pearson *r* | BH-adjusted *P* |
|---|---|---|---|
| Colonic methylation | Overall cognition score (Median-RankCombinedScore) | 0.589 | 9.87 × 10<sup>-7</sup> |
| Colonic methylation | Reciprocal of probe-trial latency | 0.526 | 2.30 × 10<sup>-5</sup> |
| Colonic methylation | Reciprocal of retention-test latency | 0.461 | 3.82 × 10<sup>-4</sup> |
| Colonic methylation | Intercept of the logarithmic regression of training latency | 0.367 | 0.0104 |
| Colonic methylation | Intercept of the regression of training cognition scores | 0.359 | 0.0104 |
| Colonic methylation | Retention cognition score | 0.356 | 0.0104 |
| Hippocampal methylation | Slope of the logarithmic regres-sion of training latency | 0.307 | 0.0380 |

To assess whether these associations persisted after accounting for age-group differences, we computed the age-adjusted correlation coefficient, *ρ*_5*jk*|age_. Nevertheless, we could not find any statistically significant 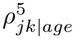. These results support an association between colonic methylation and cognition across age groups, whereas no significant age-independent association was detected.

### 2.5 Enrichment of enteric neuronal and glial markers

In spite of the tight relation between cognition score and colon methylation, enrichment analyses rarely detect enrichments related to colon. This missing enrichment of colon might be understood as follows. Although colon is full of nervous system i.e., Enteric Nervous System (ENS), since enrichment analysis using gene expression/methylation is largely overlapped with CNS, most of enrichment related to ENS can be detected as those related to CNS. To assess enrichment for enteric neuronal and glial markers among the selected genes, we performed an overlap analysis using published marker sets. Table 4 presents the neuronal, glial, and fibroblast results from Supplementary Table S4 of [11] (see Methods). Benjamini–Hochberg correction was applied jointly to 32 tests: the 29 cell-type categories from Supplementary Table S4, the neuronal and glial signature lists explicitly reported in the article’s Methods [11], and the union of those two lists. This strongly suggests the enrichment of ENS as well as that of a non-neural stromal category (Fibroblast).

**Table 4.** Enrichment of enteric neuronal, glial, and fibroblast markers among the 1,577 selected genes. Marker sets were derived from the `Mouse Colon All` worksheet of Supplementary Table S4 of Drokhlyansky et al. [11], retaining genes with `coefD` *>* 0 and padjD *<* 0.05. Overlap is reported as the number of selected marker genes divided by the number of marker genes represented in the 16,080-gene background. *P*-values were calculated using one-sided Fisher’s exact tests, and BH-adjusted *P*-values were calculated across all 32 tests described in the Methods. Fibroblasts are shown as a separate stromal category, not as a component of the enteric nervous system.

| Marker set | Overlap | Fold enrichment | *P*-value | BH-adjusted *P* |
|---|---|---|---|---|
| Enteric neuron (Neuron_1) | 38/76 | 5.10 | 5.27 × 10<sup>−19</sup> | 1.69 × 10<sup>−17</sup> |
| Enteric neuron (Neuron_2) | 15/75 | 2.04 | 5.60 × 10<sup>−3</sup> | 1.38 × 10<sup>−2</sup> |
| Enteric glia (Glia) | 31/68 | 4.65 | 2.45 × 10<sup>−14</sup> | 3.92 × 10<sup>−13</sup> |
| Fibroblast (Fibroblast) | 27/63 | 4.37 | 7.05 × 10<sup>−12</sup> | 4.51 × 10<sup>−11</sup> |

## 3 Methods

### 3.1 Gene expression profiles

Processed, variance-stabilized RNA-seq expression matrices for the hippocampus and colon were obtained from the authors’ Zenodo repository (DOI: 10.5281/zenodo.15849341). The file, ScriptsAndFilesCognitiveAgingManuscript.zip, was downloaded from https://doi.org/10.5281/zenodo.15849341 (accessed on 6th Oct 2026). By expanding the file whose structure is the directory, filesManuscript/DataAndCode/Inputs/, two files, varianceStabilizedRNASeq Brain.tsv for the brain and varianceStabilizedRNASeq Colon.tsv for colon, are extracted, respectively.

### 3.2 Methylation profiles

Processed RRBS-derived promoter methylation matrices for the hippocampus and colon were obtained from the authors’ Zenodo repository (DOI: 10.5281/zenodo.15849341). Transcript-level promoter methylation values were mapped to Ensembl gene IDs and averaged across transcripts belonging to the same gene. Two files, brainMeanPromoterMethylationNoMissingValues.tsv for brain and intestineMeanPro-moterMethylationNoMissingValues.tsv for colon, were extracted from the same zip file, ScriptsAndFilesCognitiveAgingManuscript.zip, as that of gene expression profiles.

### 3.3 Cognition score

Cognitive scores were obtained from the authors’ publicly available data archive, ScriptsAndFilesCognitiveAgingManuscript.zip, deposited in Zenodo (doi:10.5281/zenodo.15849341). Specifically, the MedianRankCombinedScore column in filesManuscript/DataAndCode/Inputs/df allCognitionData 2025-02-17.tsv was used as the cognition score. These author-provided scores were used directly without recalculation. Behavioral records were matched to the hippocampal and colonic omics samples using individual mouse identifiers after removing leading and trailing whites-pace. The analysis included the 75 mice used to construct the tensor, comprising 15 mice from each of the five age groups.

### 3.4 Tensor formation

Sample counts in the original public datasets are shown in Table 5. After matching individual identifiers across datasets, applying the exclusion criteria, and selecting 15 individuals per age group without replacement, 75 unique mice were retained for analysis (Table 6).

**Table 5.** Sample counts in the publicly available molecular data matrices and behavioral dataset before intersection, exclusion, and selection for tensor construction. Behavioral counts include individuals with missing values in one or more behavioral measures. Age-group labels follow the original sample identifiers; 2M, 9M, 15M, 24M, and 30M correspond to reported ages of 3, 9, 15, 24, and 28 months, respectively. The subsequent selection of 75 unique individuals is summarized in Table 6.

| Data matrix or dataset | 2M | 9M | 15M | 24M | 30M | Total |
|---|---|---|---|---|---|---|
| Hippocampal RNA-seq | 15 | 16 | 16 | 17 | 18 | 82 |
| Hippocampal DNA methylation | 16 | 16 | 16 | 17 | 18 | 83 |
| Colon RNA-seq | 16 | 16 | 16 | 17 | 18 | 83 |
| Colon DNA methylation | 16 | 16 | 16 | 17 | 18 | 83 |
| Behavioral dataset | 16 | 16 | 16 | 17 | 18 | 83 |

**Table 6.**
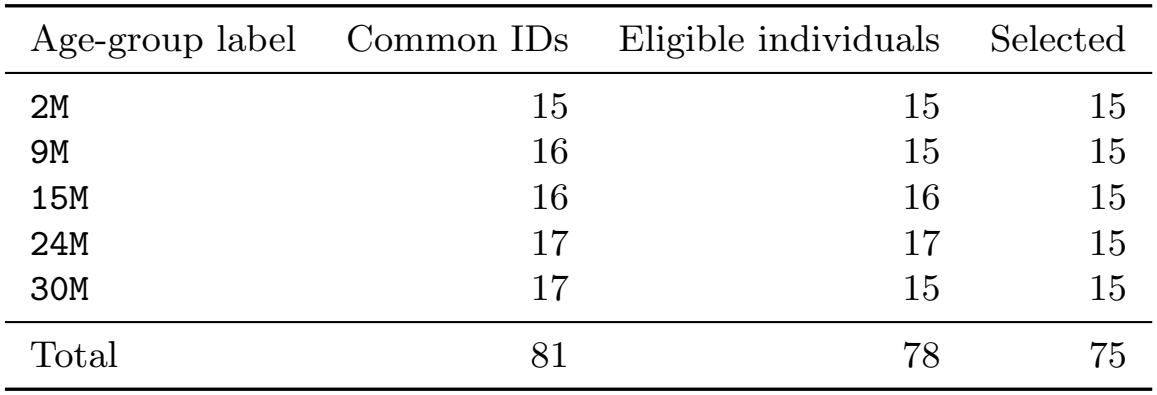
Sample availability and selection for tensor construction. Common IDs denote individuals represented in all four molecular data matrices and the behavioral dataset after trimming leading and trailing whitespace from identifiers. Eligible individuals had complete data for all nine behavioral measures and excluded `30M 16N`, as specified in the analysis code. Fifteen individuals per age group were selected without replacement, yielding 75 unique mice. Age-group labels follow the original sample identifiers, with the same age correspondence as in Table 5. The individual mouse identifiers are provided in Additional file 3.

| Age-group label | Common IDs | Eligible individuals | Selected |
|---|---|---|---|
| 2M | 15 | 15 | 15 |
| 9M | 16 | 15 | 15 |
| 15M | 16 | 16 | 15 |
| 24M | 17 | 17 | 15 |
| 30M | 17 | 15 | 15 |
| Total | 81 | 78 | 75 |

Before tensor construction, each molecular profile was standardized across the 16,080 retained common genes, separately for each tissue, modality, and individual. For each profile, the mean across genes was subtracted and the result was divided by the sample standard deviation across genes. Thus, each profile had zero mean and unit sample variance across genes. For DNA methylation, transcript-level promoter values were first averaged within each gene, and standardization was performed after selecting the common genes.

Specifically, the tensor entries were defined as

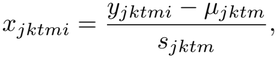

where *y*_*jktmi*_ denotes the value before standardization,

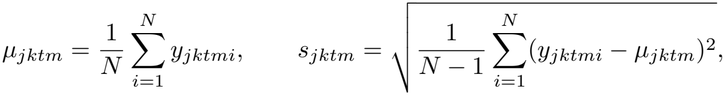

and *N* = 16,080.

The source study reports five age groups of 3, 9, 15, 24, and 28 months [10]; this age specification is also documented in GEO accession GSE302188. In the processed input files and analysis code, the corresponding sample-ID prefixes are `2M, 9M, 15M, 24M`, and `30M`, respectively. These prefixes were retained as identifiers and are not interpreted as the actual ages in months. The correspondence is recorded in Additional file 3. The profiles were formatted as a tensor of tissue (hippocampus vs colon) × modality (RNA vs methylation) × age group (3, 9, 15, 24, and 28 months) × replicate (15) × gene, *x*_*jktmi*_ ∈ ℝ^2*×*2*×*5*×*15*×N*^ where *N* = 16, 080 is the number of common genes between RNA and methylation. Within each age group, 15 eligible mice were selected without replacement and assigned to replicate positions. The same mouse occupied the same position across the four molecular profiles within an age group; replicate positions across different age groups do not denote longitudinal observations of one mouse. The saved sample-ID map is supplied in Additional file 3 and fixes the selected individuals and their ordering for reproduction.

### 3.5 Tensor decomposition

Higher order singular value decomposition (HOSVD) [9] was applied to *x*_*jktmi*_ and we got

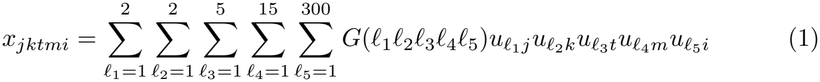

where 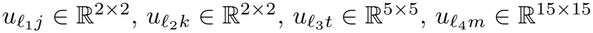, and 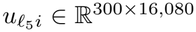, are singular value matrices. The first four factor matrices are orthogonal, whereas the gene-mode factor matrix has orthonormal rows. *G* ∈ R^2*×*2*×*5*×*15*×*300^ is the core matrix that represents the contribution of 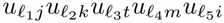 toward *x*_*jktmi*_. The reason why we restricted only top 300 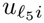 is because the number of independent singular value vectors is at most as the same as the product of the dimensions of the other four modes, 2 × 2 × 5 × 15 = 300

### 3.6 TD based unsupervised FE

In order to select critical genes, we applied TD based unsupervised FE [9] to this data set. After identification of 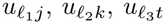, and 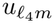 with desired properties as 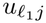 and 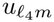 should be constant, since we seek genes whose expression is common between brain and gut as well as independent of biological replicates, 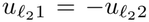 since gene expression should be suppressed with excess of methylation, and 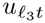 should be monotonic since the purpose of the original study is to figure out the change of cognition level upon aging. Once we get the favorite *ℓ*_1_, *ℓ*_2_, *ℓ*_3_, and *ℓ*_4_, we can investigate *G*(*ℓ*_1_*ℓ*_2_*ℓ*_3_*ℓ*_4_*ℓ*_5_) and find which *ℓ*_5_ is associated with the absolute largest value of *G*. Then using the selected 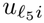, we try to select genes, *i*s.

To do this, we assume that 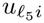 obeys Gaussian distribution (null hypothesis), attribute *P*-values to genes (*i*s) and genes (*i*s) associated with the smaller *P*-values (i.e., statistically significant) are selected. *P*-values are attributed to *i*s as

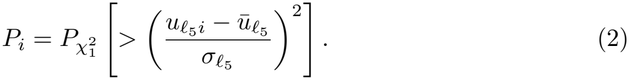

Here, 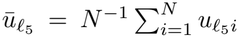, and 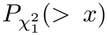 denotes the upper-tail probability of a chi-squared distribution with one degree of freedom. 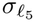 is the standard deviation of 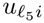.

The scale parameter 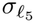 was estimated by histogram flattening using the implementation in `select_cognitive_gene_features.R`, which calls `select_features_flat_histogram`. The full-vector mean 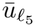 was subtracted once and held fixed throughout the search. Let *s*_0_ be the sample standard deviation of the centered loadings, computed with denominator *N* − 1. The default search interval was [0.01*s*_0_, 100*s*_0_].

For every candidate *σ, P*_*i*_(*σ*) was calculated for all *N* = 16,080 genes using the chi-squared expression above, followed by BH adjustment across all *N* genes. The candidate background was

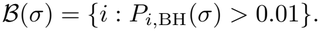

Candidates were admissible only if |*ℬ*(*σ*)| ≥ max{2, ⌈0.5*N* ⌉}; thus at least 8,040 background genes were required. For admissible candidates, a histogram of 1 − *P*_*i*_(*σ*) for *i* ∈ *ℬ*(*σ*) was constructed using 100 equal-width bins on [0, 1]. Bins were right-closed, with zero included in the first bin. Writing *h*_*b*_(*σ*) for the count in bin *b* and 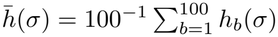, the objective was

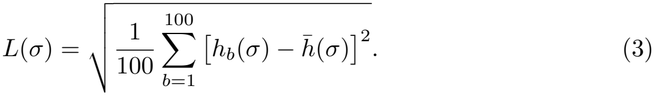

Inadmissible candidates were assigned infinite loss. This is the population standard deviation of the bin counts, as implemented by `sqrt(mean((h-mean(h))^2))`.

The coarse search used 101 equally spaced points in log *σ* over the search interval, adding *s*_0_ if it was not already on the grid. Two refinement rounds were then performed. Each round evaluated 61 equally spaced log-scale points within one current grid step on either side of the best candidate, clipped to the original search interval. The initial step was the coarse log-grid spacing; after each refinement, it was updated to that refinement’s log-grid spacing. The best candidate was chosen from all admissible evaluations accumulated so far. Ties in *L* were resolved first by proximity to *s*_0_ on the logarithmic scale and then by choosing the smaller *σ*. Accordingly, the reported optimum is the minimum among the evaluated candidates, not a claim of exact continuous global optimization.

At the chosen scale, *P*-values and BH-adjusted *P*-values were recalculated for all genes, including those excluded from the candidate background. Genes with *P*_*i*,BH_ ≤ 0.01 were selected. The saved analysis used *σ* ≃ 0.00341161 and selected 1,577 genes from the fifth gene-mode singular vector. The selector script and an explicit parameter summary are included in Additional file 3; the full workflow is available from the repository specified in the Declarations.

### 3.7 Enrichment analysis

To see whether the selected genes are associated with desired biological functions/meaning, the selected genes are uploaded to Enrichr [12]. All 228 libraries returned by the Enrichr library catalog at the time of analysis were queried using 1,574 unique gene symbols. The standard Enrichr background was used; the experimental 16,080-gene background was not supplied to this service. The adjusted *P*-values returned by Enrichr apply within each library and were not corrected jointly across all 228 libraries. No separate mouse-to-human ortholog conversion was performed. Terms relevant to aging and neural functions were highlighted for interpretation. Additional file 2 contains the archived results and input list; this analysis is distinct from the measured-background marker tests and the local Boruta-excluded analysis described below.

### 3.8 Relating *u*_**5*i***_ to cognition scores

First, gene expression as well as methylation profiles of individual mice, *x*_*jktmi*_ were projected to 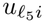 as

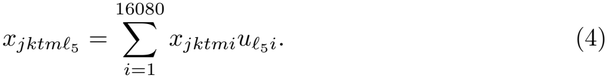

Two-sided Pearson correlation tests were performed across the 75 unique mice for each of the four molecular profiles and nine behavioral measures (36 tests). BH correction was applied jointly to these 36 unadjusted tests. The complete results are provided in Additional file 3.

The correlation is also computed with excluding age effect as

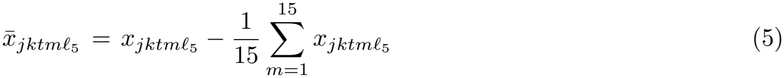

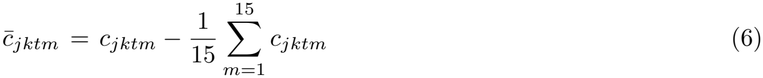

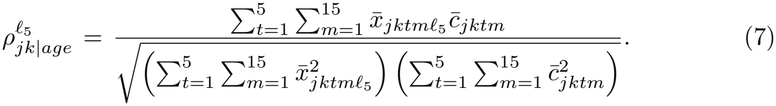

For age adjustment, the molecular and behavioral scores were each residualized against an intercept and a categorical five-level age factor, equivalent to subtracting the respective age-group means as above. The correlation of these residuals was tested using 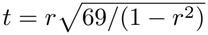 with 69 degrees of freedom (75−5−1), using a two-sided test. BH correction was applied separately across the 36 age-adjusted tests; all adjusted *P*-values exceeded 0.05. These full results are also included in Additional file 3.

### 3.9 Enrichment of enteric neuronal and glial markers

To assess whether the selected genes were enriched for enteric neuronal and glial markers, we evaluated their overlap with published marker sets using one-sided Fisher’s exact tests.

Marker sets were constructed from Supplementary Table S4 of Drokhlyansky et al. [11], using the Mouse Colon All worksheet. For each cell-type category, we retained the reported genes with a positive coefficient for the discrete component of the MAST model (coefD *>* 0) and an adjusted *P*-value below 0.05 (padjD *<* 0.05). These comparisons contrast each cell category with all remaining cells in the reference dataset. The same criteria were applied to all 29 categories, without further restricting the number of markers or selecting markers based on their overlap with our selected genes. The resulting sets therefore represent filtered markers reported in the supplementary table, rather than an exhaustive reconstruction of markers from the raw single-cell data.

The original category labels were retained. In particular, `Neuron_1` and `Neuron_2` were treated as separate enteric neuronal marker sets, and `Glia` as an enteric glial marker set. The `Fibroblast` category was evaluated separately as a non-neural stromal category.

Marker symbols were mapped to mouse Ensembl gene identifiers using the annotation supplied with the original cognitive-aging dataset and restricted to the 16,080 genes included in our tensor analysis. Ensembl gene identifiers were used as the counting unit; when a symbol corresponded to multiple background gene identifiers, each identifier was retained. The selected set comprised 1,577 genes, all contained in this background.

Enrichment was assessed using one-sided Fisher’s exact tests with the alternative hypothesis of overrepresentation. Fold enrichment was calculated as

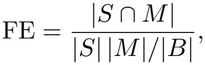

where *S* denotes the selected genes, *M* the marker genes represented in the background, and *B* the background gene set. BH correction was applied jointly to all 32 marker-set tests described in the Results. The full tests, marker membership, and background annotation are provided in Additional file 3.

### 3.10 Enrichment analysis after exclusion of Boruta-selected genes

Enrichment was recalculated locally using gene sets downloaded from two Enrichr libraries: `GO_Biological_Process_2026` for biological processes and `PanglaoDB_Augmented_2021` for cell-type signatures. The synaptic terms in Table 7 were obtained from the former library, whereas “Enteric Neurons” and “Enteric Glia Cells” were obtained from the latter. Gene symbols were converted to uppercase and deduplicated before matching to the library gene sets.

**Table 7.**
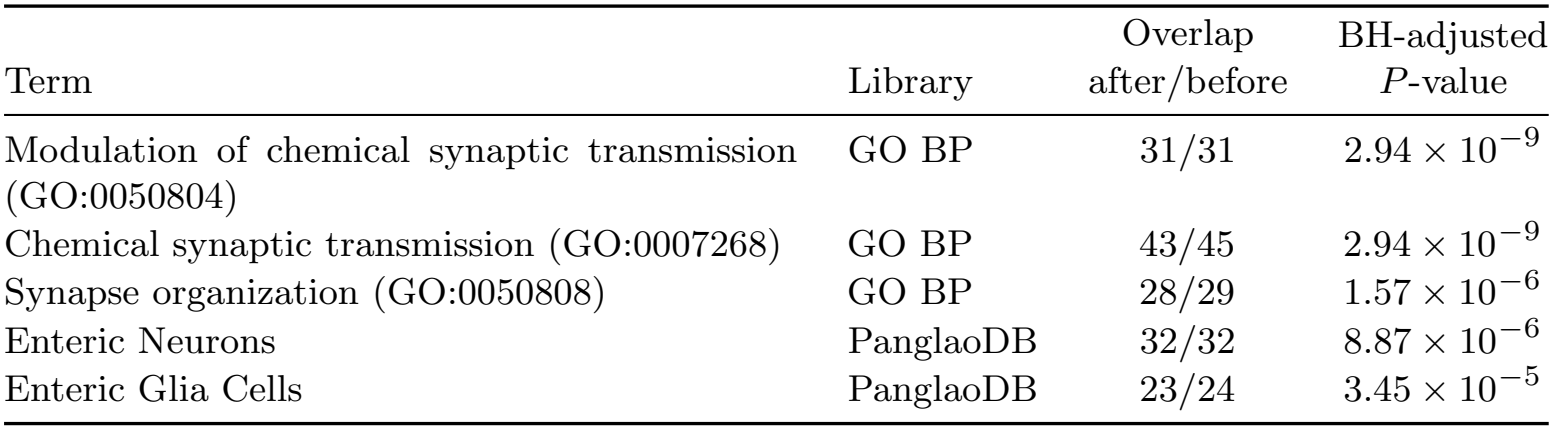
Selected synaptic and enteric cell-type enrichment results after exclusion of Boruta-selected genes. GO terms were obtained from `GO_Biological_Process_2026` (GO BP), and cell-type signatures from `PanglaoDB_Augmented_2021` (PanglaoDB). The overlap column reports the number of term-associated tensor-selected genes remaining after Boruta exclusion relative to the corresponding number before exclusion; the denominator is not the total size of the term’s gene set. Adjusted *P*-values were calculated from upper-tail hypergeometric tests using library-specific backgrounds after Boruta exclusion, with Benjamini–Hochberg correction across all terms in each library.

| Term | Library | Overlap<br>after/before | BH-adjusted<br>*P*-value |
|---|---|---|---|
| Modulation of chemical synaptic transmission (GO:0050804) | GO BP | 31/31 | 2.94 × 10<sup>−9</sup> |
| Chemical synaptic transmission (GO:0007268) | GO BP | 43/45 | 2.94 × 10<sup>−9</sup> |
| Synapse organization (GO:0050808) | GO BP | 28/29 | 1.57 × 10<sup>−6</sup> |
| Enteric Neurons | PanglaoDB | 32/32 | 8.87 × 10<sup>−6</sup> |
| Enteric Glia Cells | PanglaoDB | 23/24 | 3.45 × 10<sup>−5</sup> |

For each library *L*, let *A*_*L*_ denote the union of genes annotated in that library and *B* the Boruta exclusion set, both represented by gene symbols. The background was defined as

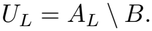

The query set was defined as *Q*_*L*_ = *T* ∩ *U*_*L*_, where *T* denotes the symbols of the 1,513 remaining tensor-selected genes. Each term-specific gene set *G*_*Lj*_ was restricted to this background as *M*_*Lj*_ = *G*_*Lj*_ ∩ *U*_*L*_. Thus, the background consisted of library-annotated genes after Boruta exclusion, rather than all genes measured in the experiment.

Overrepresentation was evaluated using the upper-tail hypergeometric test, equivalent to a one-sided Fisher’s exact test with fixed margins. For *N* = |*U*_*L*_|, *n* = |*Q*_*L*_|, *K* = |*M*_*Lj*_|, and *k* = |*Q*_*L*_ ∩ *M*_*Lj*_|, the enrichment *P*-value was

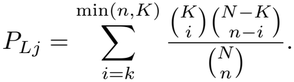

The GO library contained 15,287 background genes and 730 query genes, whereas the PanglaoDB library contained 6,479 background genes and 525 query genes. Benjamini– Hochberg correction was applied separately within each library across all tested terms: 5,954 GO terms and 178 PanglaoDB cell-type signatures. The adjusted *P*-values were therefore recalculated locally and were not taken from the original Enrichr output.

The full 6,132 term-level tests, library-specific settings, Boruta feature mappings, and the 1,513 remaining genes are supplied in Additional file 3.

### 3.11 Generative AI assistance

ChatGPT (OpenAI) was used to assist with code preparation, data-processing checks, and manuscript editing. The authors are responsible for the analyses, interpretation, and final content of the manuscript.

## 4 Discussion

Although TD based unsupervised FE applied to brain and colon gene expression and methylation profiles successfully select the set of genes that might be critical to brain-gut axis for cognition with aging, if the results are identical to those in the original research [10], it is not so promising. Thus it is important to identify which is common and which is not. Flor et al [10] also detected the significant correlation between cognition scores and colon methylation but not brain gene expression/methylation, colon/gene expression. Thus the primary findings are common between Flor et al [10] and us. The primary difference is as follows. At first, the number of detected genes is much larger in our study. We identified 1,577 genes, compared with 506 genes selected in at least one individual or integrated Boruta analysis by Flor et al. [10], including analyses of liver data, which were not included in our study. The overlap between these 506 genes and our 1577 genes is as small as 64. If restricted to those detected at least once in Boruta integrated study, the number of genes decreases to 182, of which only 31 overlap with our 1,577 genes. When restricted to those detected at least ten times in Boruta integrated study, the number of genes decreases further to 157, which has only 26 overlaps with our 1577 genes. Thus, ours are not only larger than Flor et al [10], but also highly distinct.

With enrichment analysis after exclusion of Boruta-selected genes, we can show that ours have enough significant enrichment even excluding them. To identify functional associations among genes selected by unsupervised tensor decomposition but not by Boruta using cognitive scores, we excluded the broadest available Boruta gene set. This exclusion set comprised the union of genes corresponding to features selected in at least one Boruta run across all analyzed feature sets, including the combined feature set. Transcript identifiers were mapped to gene identifiers using the annotation supplied with the dataset. Of the 1,577 tensor-selected genes, 1,513 remained after exclusion.

Synaptic processes and enteric neuronal and glial signatures remained significantly enriched after exclusion of Boruta-selected genes (Table 7). All 32 genes overlapping the Enteric Neurons signature were retained, including *ELAVL4, HAND2, STMN2, SLC18A3, SLC5A7, NOVA1*, and *RBFOX1*. These results indicate that unsupervised tensor decomposition identified genes associated with synaptic functions and enteric cell-type signatures that were not selected by Boruta using cognitive scores in the analyzed runs. This enrichment is conditional on the library-specific backgrounds and does not by itself establish tissue of origin or a causal relationship with cognitive function.

Our analysis recovered the reported association between colonic methylation and cognition without using cognitive scores for gene selection. It also identified a larger, largely distinct gene set that retained significant enrichment for synaptic processes and enteric cell-type signatures after exclusion of Boruta-selected genes.

## 5 Conclusions

Tensor decomposition-based unsupervised feature extraction identified a shared age-related molecular component in hippocampal and colonic transcriptomic and DNA methylation profiles without using cognitive scores for gene selection. The association between the resulting colonic methylation score and cognition was consistent with the principal relationship reported in the original supervised analysis. The absence of significant associations after age adjustment indicates that the present evidence concerns variation across age groups rather than an established relationship with individual cognitive differences independent of age.

The contribution of this analysis extends beyond recovering that association. Of the 1,577 selected genes, 1,513 were absent from the broadest available Boruta-selected set, yet retained enrichment for synaptic processes and enteric neuronal and glial signatures. Together with enrichment of independently defined enteric cell-type markers in the full selected set, these results show that the additional genes form biologically coherent groups relevant to investigating the gut–brain axis.

Our findings support unsupervised tensor integration as a complementary strategy for identifying candidate genes that are not recovered by cognitive-score-guided feature selection. They do not establish that these genes originate specifically from enteric cells or mediate a causal influence of the gut on cognition. Independent cohorts and cell-type-resolved functional studies will be needed to determine how the identified signatures relate to enteric neural changes and cognitive aging.

### Supplementary information. Additional file 1

File: `Additional file 1.csv` (CSV). Title: Selected genes and gene-symbol mappings. Description: The 1,577 selected Ensembl mouse gene identifiers, corresponding gene symbols, and mapping status. These genes correspond to 1,574 unique symbols after deduplication.

### Additional file 2

File: `Additional_file_2.zip` (ZIP). Title: Archived Enrichr results across 228 libraries. Description: All available archived term-level exports for the 228 queried libraries, input symbols, library summaries, significant-term summaries, analysis metadata, scripts, and an English README describing backgrounds and multiple testing.

### Additional file 3

File: `Additional_file_3.zip` (ZIP). Title: Complete statistical results and analysis reference files. Description: All 36 unadjusted and 36 age-adjusted correlation tests; all 32 marker-enrichment tests; all 6,132 Boruta-excluded enrichment tests; marker membership and background annotation; Boruta feature mappings and overlap summaries; the 1,513 tensor-only genes; individual mouse identifiers, molecular scores, and matched behavioral measures; the age-label correspondence; and the histogram-based selector script and parameter summary. An English README explains the files and their provenance.

## Declarations

### Ethics approval and consent to participate

Not applicable to this secondary analysis of publicly available data. No new animal or human experiments were performed. The source mouse datasets were generated in the study of Flor et al. [10].

### Consent for publication

Not applicable.

### Availability of data and materials

The original processed molecular and behavioral data are available from Zenodo at https://doi.org/10.5281/zenodo.15849341. The selected genes, archived enrichment outputs, and complete downstream statistical results and supporting reference files are provided in Additional files 1–3. R code for the analysis is available at https://github.com/tagtag/TDbasedUFE_Brain-Colon. The selector used to document the histogram optimization is also included in Additional file 3.

### Authors’ contributions

Y.-h.T. conceived and planned the study, curated the data, and performed the analyses. Y.-h.T. and T.T. evaluated the results and wrote and reviewed the manuscript. Both authors read and approved the final manuscript.

## Acknowledgements

Not applicable.

